# Immunological imprinting of humoral immunity to SARS-CoV-2 in children

**DOI:** 10.1101/2022.07.26.501570

**Authors:** Alexander C. Dowell, Tara Lancaster, Rachel Bruton, Georgina Ireland, Christopher Bentley, Panagiota Sylla, Jianmin Zuo, Sam Scott, Azar Jadir, Jusnara Begum, Thomas Roberts, Christine Stephens, Shabana Ditta, Rebecca Shepherdson, Annabel A. Powell, Andrew J. Brent, Bernadette Brent, Frances Baawuah, Ifeanyichukwu Okike, Joanne Beckmann, Shazaad Ahmad, Felicity Aiano, Joanna Garstang, Mary E. Ramsay, Rafaq Azad, Dagmar Waiblinger, Brian Willett, John Wright, Shamez N. Ladhani, Paul Moss

## Abstract

Omicron variants of SARS-CoV-2 are globally dominant and infection rates are very high in children. We determined immune responses following Omicron BA.1/2 infection in children aged 6-14 years and related this to prior and subsequent SARS-CoV-2 infection or vaccination. Primary Omicron infection elicited a weak antibody response with poor functional neutralizing antibodies. Subsequent Omicron reinfection or COVID-19 vaccination elicited increased antibody titres with broad neutralisation of Omicron subvariants. Prior pre-Omicron SARS-CoV-2 virus infection or vaccination primed for robust antibody responses following Omicron infection but these remained primarily focussed against ancestral variants. Primary Omicron infection thus elicits a weak antibody response in children which is boosted after reinfection or vaccination. Cellular responses were robust and broadly equivalent in all groups, providing protection against severe disease irrespective of SARS-CoV-2 variant. Immunological imprinting is likely to act as an important determinant of long-term humoral immunity, the future clinical importance of which is unknown.

## Introduction

The emergence of the SARS-CoV-2 Omicron variant (B.1.1.529) marked a major shift in the COVID-19 pandemic. The acquisition of multiple mutations, primarily within the spike receptor binding domain (RBD), enhanced Omicron infectivity and led to substantial evasion of antibody responses elicited from prior infection or vaccination (*1*–*3*). As such, the variant demonstrated remarkable capacity to outcompete pre-Omicron SARS-CoV-2 and became the dominant variant globally.

Omicron infection rates have been high in all age groups irrespective of prior infection or vaccination status but have been particularly notable in children (*4*). Reinfection with Omicron subvariants is also observed (*5*) and protection against reinfection appears lower than was seen in children following infection with pre-Omicron SARS-CoV-2 (*6*). Studies of children who have been infected with pre-Omicron SARS-CoV-2 virus or have undergone vaccination reveals relatively low antibody binding to Omicron, in a similar pattern to that seen in adults (*1*, *7*–*9*). Less is known regarding the nature of the immune response elicited by primary Omicron infection in this age group, although low antibody responses have been reported (12) and concur with similar findings in adults (*10*, *11*). Prior infection status with pre-Omicron variants is also suggested to be an important determinant of response to Omicron in adults (*7*, *11*, *12*).

Profound differences are observed in the age-dependency of the immune response against SARS-CoV-2. Infections are generally mild or asymptomatic in children and potential determinants of this may include increased expression of IFN-response genes within the mucosal epithelium and enhanced innate immune responses (*13*, *14*). Reassuringly, children develop robust systemic adaptive immune responses against pre-Omicron variants, even after asymptomatic infection (*15*), with increased clonality compared to adults (*14*), although it is not known if similar immune responses will be observed after Omicron infection.

Here we studied antibody and cellular immunity in children following a recent Omicron infection. Furthermore, we related immune response to prior infection or vaccination and also assessed the subsequent impact of reinfection or vaccination. Primary Omicron infection was weakly immunogenic whilst the profile of humoral response was markedly influenced by prior infection or vaccination and matured substantially following reinfection. Cellular immunity was robust in all cases. These data have important implications for understanding the immune responses against SARS-CoV-2 in children and guiding optimal immune protection.

## RESULTS

### Primary Omicron infection in children elicits low antibody titre with poor neutralisation activity

Blood samples from the sKIDS (n=29) and Born in Bradford (n=14) cohort studies were available from 43 unvaccinated children who had recently tested positive for Omicron BA.1/2 infection (**Extended Data Figure 1**; **Extended Data Table 1**). The sKIDs study of children aged 6-14 years (*16*) and the Born in Bradford (BiB) study of children aged 8-15 years are blood sampling studies of children during the COVID-19 pandemic and their design and sampling dates are explained fully in the Methods section (*17*).

Initial studies were undertaken to assess how antibody responses following Omicron compared in children in whom this represented their primary SARS-CoV-2 infection or those who had previously had a pre-Omicron infection. Serological samples prior to the Omicron infection were available from 15 children of whom 4 were SARS-CoV-2 seronegative (primary infection) and 11 seropositive (secondary infection). Primary Omicron infection elicited low antibody titres against both the ancestral spike and RBD domains (**Figure 1A, Extended Data Figure 2A**), consistent with previous reports (*18*). Titres against Delta variant were high in one child, likely due to subclinical Delta infection between blood sampling, and as such this was regarded as a secondary infection (**Extended Data Figure 2A**). In contrast to the profile of primary infection, all secondary Omicron infections elicited a substantially higher antibody response.

**Figure 1.**
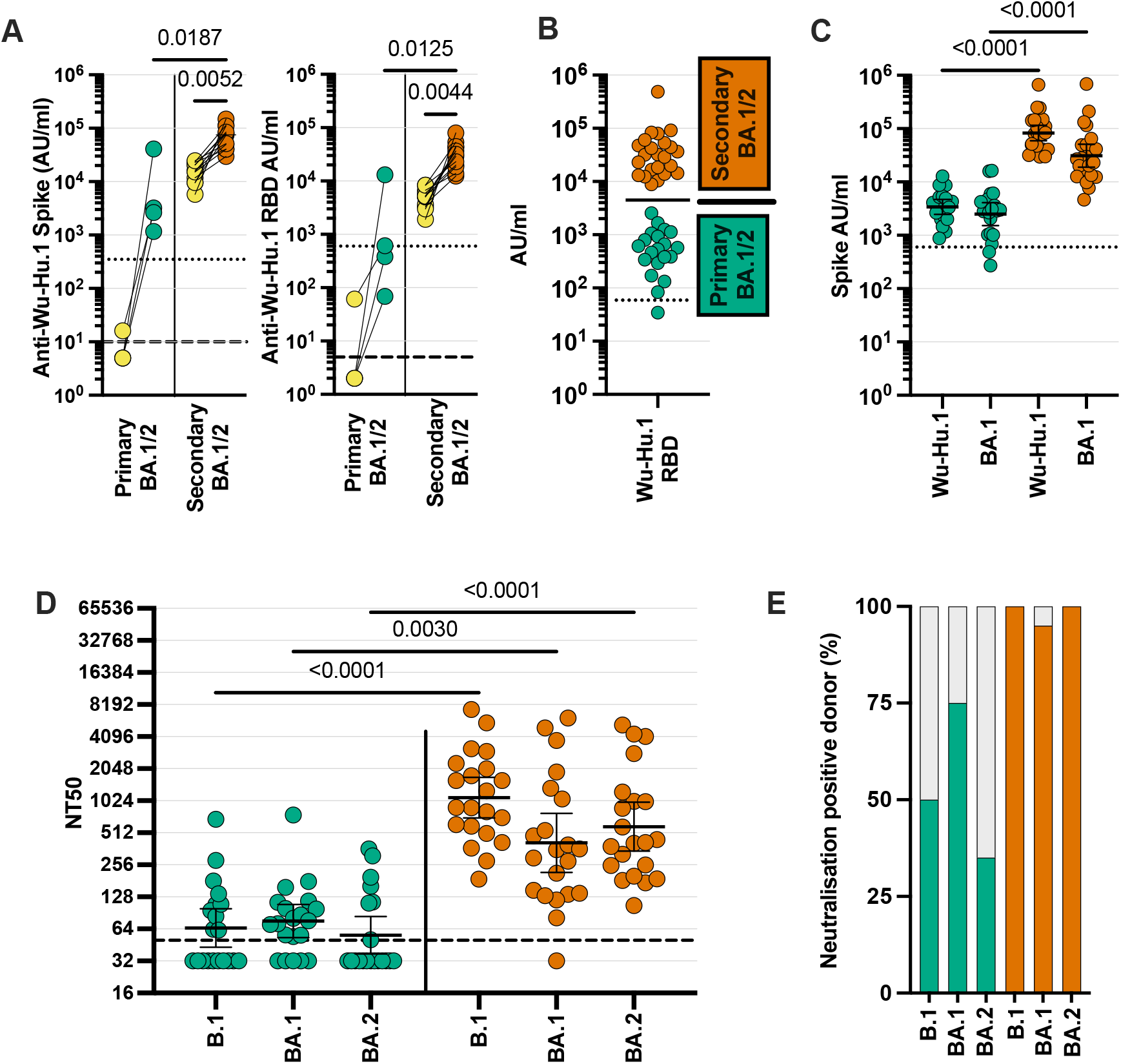
Low titre antibodies and neutralisation develop following primary Omicron infection. SARS-CoV-2-specific antibody responses were determined in children with recent Omicron BA.1/2 infection. **A**. Wuhan-Hu-1 (Wu-Hu-1) spike antibody binding in samples from children who were SARS-CoV-2 seronegative (n=4) or seropositive (n=11) prior to Omicron infection (yellow). Dotted lines indicate seropositive cut-off for Wuhan-Hu-1-specific antibody response; dashed lines indicate below limit of detection. Friedman Test with Dunn’s multiple comparisons test. **B**. Wuhan-Hu-1 RBD-specific antibody binding in the total cohort of 43 children with recent Omicron BA.1/2 infection shows bimodal distribution of primary (green; n=20) or secondary (orange; n=23) SARS-CoV-2 infection status. Bar indicates geometric mean; representative of primary or secondary omicron infection. **C.** Wuhan-Hu-1 and Omicron BA.1 spike-specific antibody binding in children with primary (green; n=20) or secondary (orange; n=23) Omicron BA.1/2 infection. Dotted line indicates seropositive cut-offs for Wuhan Hu-1. Friedman Test with Dunn’s multiple comparisons test, bars indicate geometric mean ±95% CI. **D**. Neutralisation of pseudovirus bearing B.1 (Wuhan-Hu-1, D614G) ancestral or Omicron BA.1 or BA.2 sequence spike protein in children following primary (green; n=20) or secondary (orange; n=21) Omicron infection. Dashed lines indicate limit of detection. Kruskal-Wallis test with Dunn’s multiple comparisons test, bars indicate geometric mean ±95% CI. **E**. Bar graph indicating the percentage of children with detectable neutralizing titres to B.1 or Omicron BA.1 or BA.2 following primary (n=20) or secondary (n=21) Omicron infection.

This bimodal profile of antibody response was also apparent across the total cohort of 43 children with recent Omicron infection and was therefore used to define primary (n=20) and secondary (n=23) infection (**Figure 1B, Extended Data Figure 2B**). Due to the cross-sectional nature of the collection, samples were taken at different timepoints following infection but this did not impact on antibody titre (**Extended Data Figure 3A)**.

The magnitude and relative antibody response to both the ancestral or Omicron BA.1 infection was then determined following primary or secondary infection (**Figure 1C, Extended Data Figure 4**). Primary Omicron infection elicited broadly equivalent antibody responses against all variants although these were of low titre. Notably, Omicron-specific titres did not exceed those seen in historical samples following infection with a pre-Omicron variant (**Extended Data Figure 4**). In contrast, antibody responses after secondary Omicron infection were 24-fold and 12-fold higher against ancestral and Omicron BA.1 spike (**Extended Data Table 2**).

Functional humoral immunity was assessed with a pseudovirus-based neutralisation assay following primary (n=20) or secondary (n=21) Omicron infection. Neutralisation capacity was low following primary infection and it was notable that 25% and 65% of children did not have a measurable neutralisation titre against BA.1 or BA.2 respectively. In contrast, neutralisation was robust following secondary infection (**Figure 1D, E**).

These data show that primary Omicron infection in children elicits a low titre antibody response with poor neutralising activity. In contrast, prior infection with a pre-Omicron virus primes the immune system to develop robust humoral immunity to all viral variants following Omicron infection.

### Re-infection or COVID-19 vaccination markedly boosts antibody responses following primary Omicron infection

The endemic human coronaviruses OC43, HKU-1, 229E and NL63 (HCoV) are common aetiological agents of upper respiratory tract infection in children and repeated infection acts to incrementally elevate virus-specific antibody titres (*19*). As such, we next assessed whether an Omicron reinfection could similarly act to boost humoral immunity following primary Omicron infection. The serological response to COVID-19 vaccination following primary infection was also determined.

Prospective follow-up serum samples were obtained from 24 unvaccinated children with prior serological evidence of primary Omicron infection. Omicron reinfection was identified on the basis of a >2-fold increase in spike-specific antibody titre, similar to previous reports of HCoV epidemiology (*20*). Reinfection was observed in 13 children (54%), consistent with reported reinfection rates following primary Omicron infection (*5*). Spike-specific antibody titres increased markedly following Omicron reinfection (>0.05, Kruskal Wallis test, with Dunn’s correction) with titres against BA.1 increasing 10-fold (**Extended Data Table 2**).

Samples were also obtained from 11 children who had received a single dose of BNT162b2 vaccine following primary Omicron infection. A strong boosting effect was seen with BA.1-specific titres increasing by 115-fold following vaccination (>0.0001, Kruskal Wallis test, with Dunn’s correction; **Extended Data Table 2;** Extended Data Figure 4C).

The magnitude of antibody response and relative neutralisation activity was then determined against ancestral spike and Omicron subvariants following primary Omicron (n=20), Omicron reinfection (n=13) or vaccination after primary infection (n=10). Antibody levels (**Figure 2A**) and neutralisation capacity (**Figure 2B**) were substantially enhanced against all variants following reinfection or vaccination.

**Figure 2.**
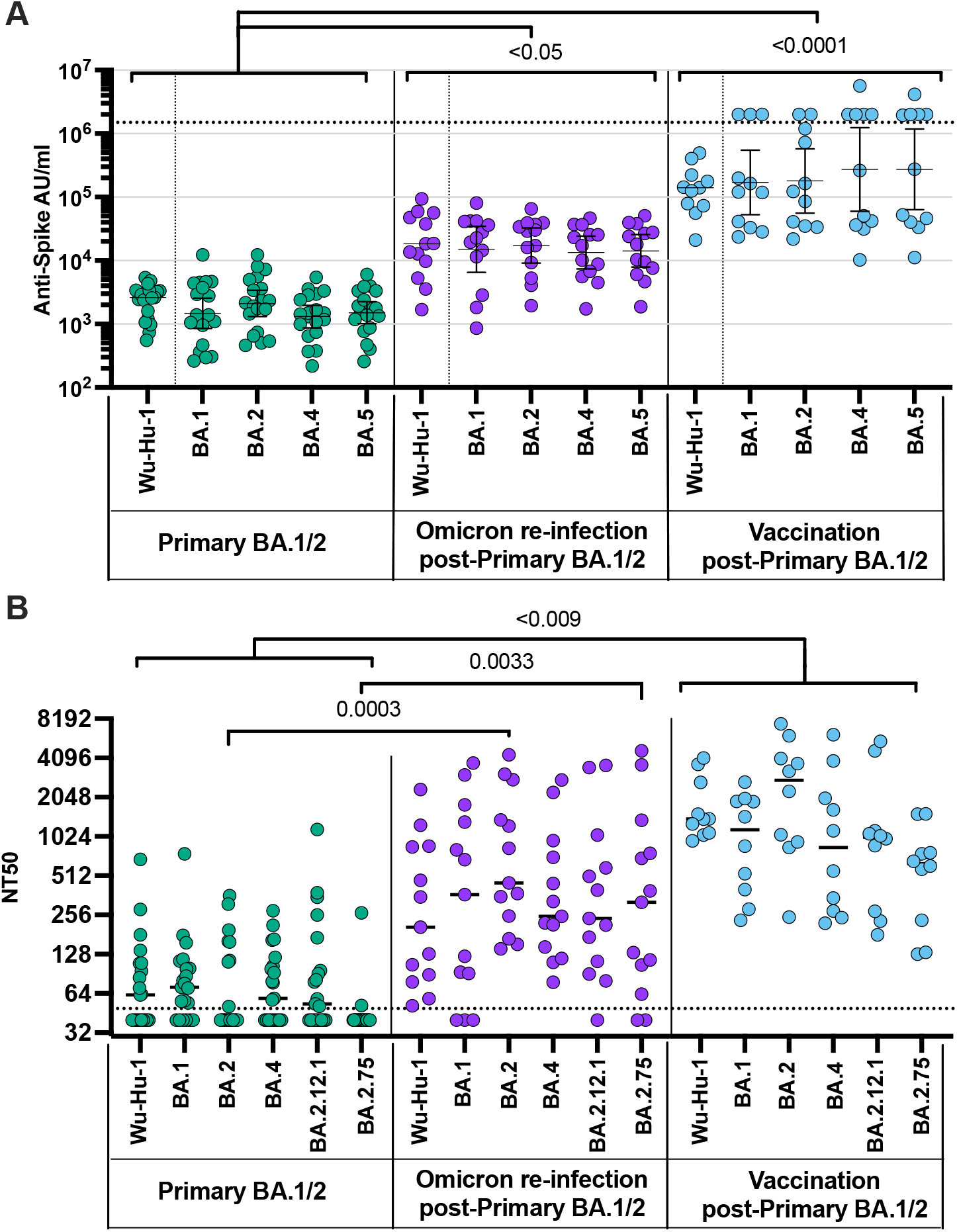
Secondary antigenic challenge enhances antibody levels and neutralisation following primary Omicron infection. (A) Antibody levels specific for spike from Wuhan-Hu-1, and Omicron BA.1, BA.2, BA.4 and BA.5 were determined in children following primary BA.1/2 Omicron infection (green; n=19), Omicron reinfection (purple; n=13) and COVID-19 vaccination (blue; n=11) after primary BA.1/2 infection. (B) Neutralisation of pseudovirus-bearing ancestral B.1 or Omicron variant spike protein in children following primary Omicron BA.1/2 infection (green; n=19), Omicron BA.4/5 reinfection (purple; n=13) or following COVID-19 vaccination after primary BA.1/2 infection (blue; n=11). Responses following secondary Omicron BA.1/2 infection are shown for comparison (n=21). Dashed lines indicate limit of detection. Kruskal-Wallis test with Dunn’s multiple comparisons test, bars indicate geometric mean ±95% CI.

These findings indicate that a second SARS-CoV-2 antigen challenge following primary Omicron infection, either in the form of reinfection or vaccination, markedly boosts spike-specific antibody magnitude and function against current Omicron subvariants.

### Primary vaccination elicits antibodies against Omicron which are not enhanced following breakthrough Omicron infection

15 children within the cohort had received mRNA COVID-19 vaccines prior to emergence of Omicron. As the vaccine contains the Wuhan-Hu-1 spike we next assessed Omicron-specific antibody titre in this cohort. Additionally, sera were obtained from 6 children with an Omicron breakthrough infection after vaccination. 10/15 and 4/6 children, respectively, had received two BNT162b2 vaccine doses.

Vaccines were strongly immunogenic and Omicron-specific antibody responses following one or two vaccine doses were 21-fold and 40-fold higher than seen after primary natural infection. Of note, prior infection history was not known and it is possible that some children had been naturally infected prior to vaccination. Antibody responses following Omicron breakthrough infection after vaccination were next assessed and seen to be comparable to those seen in the total vaccine group with no evidence of boosting from recent infection, potentially due to a ceiling effect on antibody titre (**Figure 3A**; **Extended Data Figure 4D-E).** Matched pre-infection samples were not available from children with breakthrough infection. Neutralisation titres were also similar between the two groups and comparable to those seen following dual-Omicron infection (**Figure 3B**).

**Figure 3.**
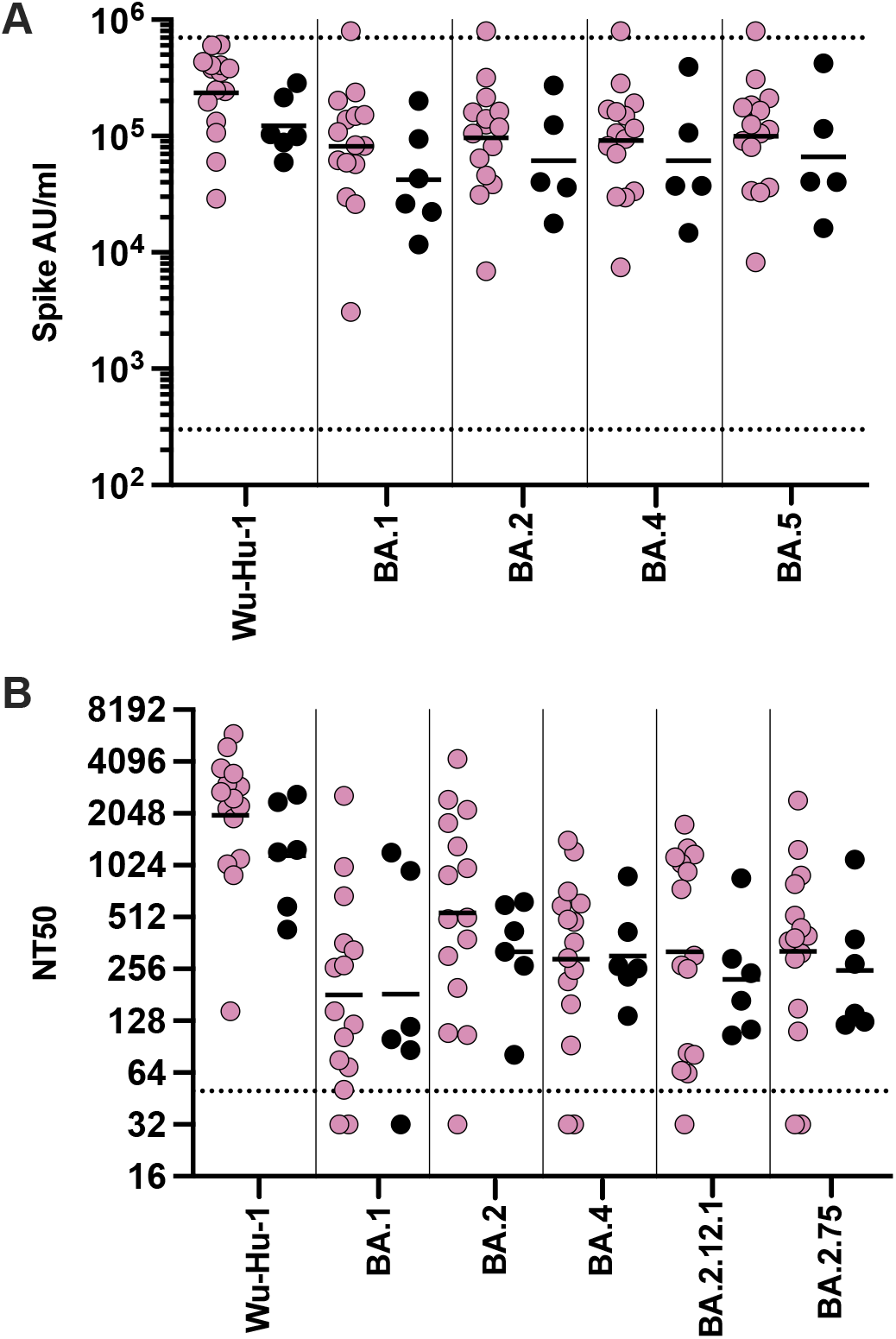
COVID-19 vaccine induces strong antibody responses that are not enhanced following breakthrough infection. (A) Antibody levels specific for spike from Wuhan-Hu-1, and Omicron BA.1, BA.2, BA.4 and BA.5 were determined in children following vaccination (pink, n=15) or BA.1/2 Omicron breakthrough infection (black, n=6) (B) Neutralisation of pseudovirus-bearing ancestral B.1 or Omicron variant spike protein in children following vaccination (pink, n=15) or BA.1/2 Omicron breakthrough infection (black, n=6). Dashed lines indicate limit of detection, bars indicate geometric mean

These data show that primary series vaccination in children induces an Omicron-specific antibody response that is higher than seen after Omicron secondary infection whilst Omicron breakthrough infection does not appear to further enhance humoral immunity.

### Initial Omicron or pre-Omicron variant challenge determines the long-term balance of antibody binding

We next determined how an initial infection with Omicron or a pre-Omicron virus acted to determine the relative balance of antibody binding against these two major viral lineages and to what extent this was modulated by subsequent challenge with Omicron re-infection or vaccination with the Wuhan-Hu-1 spike protein.

The relative ratio of antibody binding against Omicron or pre-Omicron Wuhan-Hu-1 spike was 0.7 following primary Omicron infection indicating broadly equivalent response. This value increased slightly to 0.9 following Omicron reinfection. Of note, the ratio increased to 1.3 following vaccination showing that vaccine challenge with pre-Omicron spike could not deviate antibody responses away from Omicron recognition (**Figure 4A,B**).

**Figure 4.**
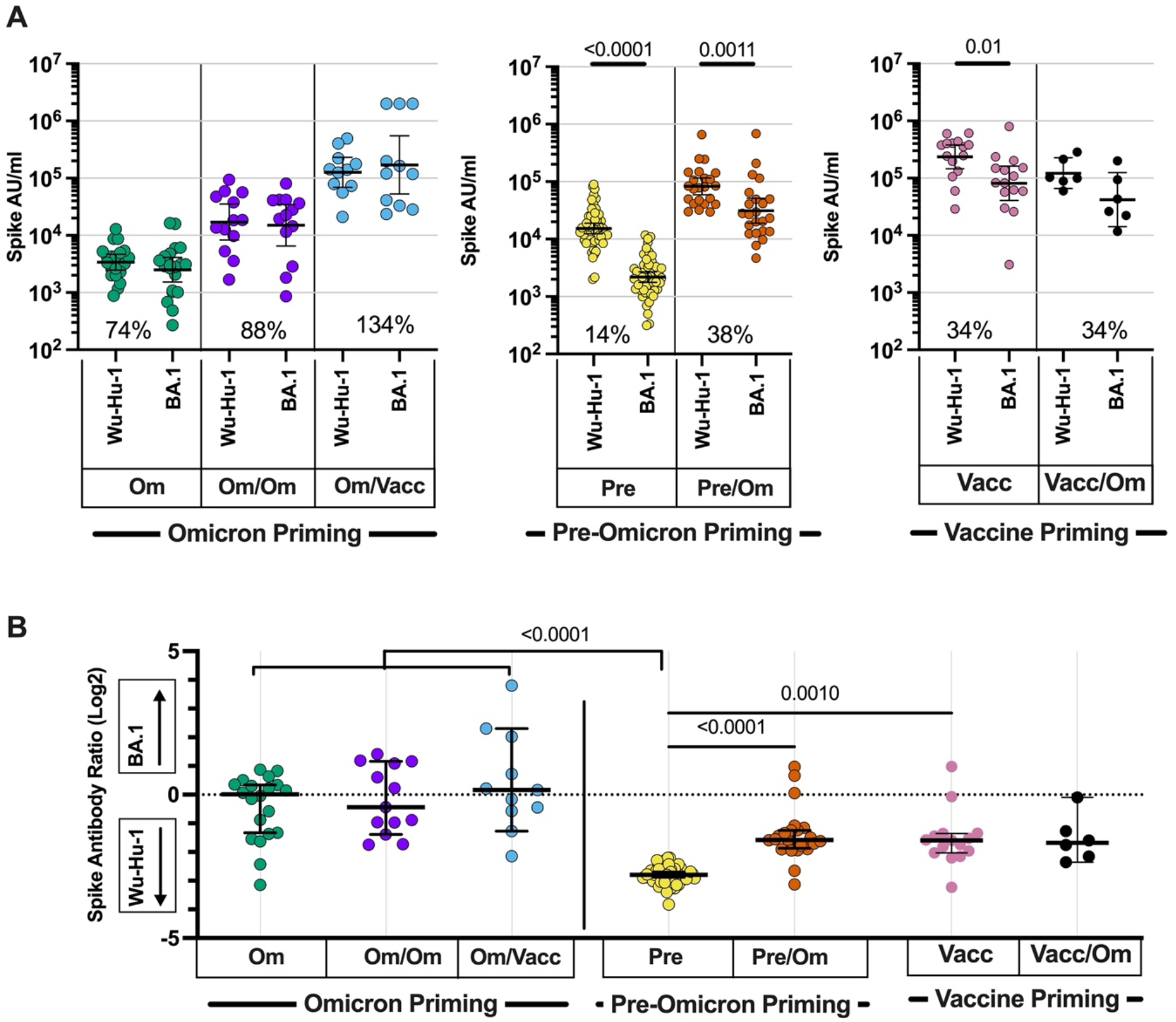
Relative antibody binding to Omicron or ancestral spike protein in relation to primary virus or vaccine challenge. A. Antibody binding against ancestral Wuhan-Hu-1 or Omicron BA.1 spike in children following three different forms of initial primary antigenic challenge. i) Initial primary Omicron infection (green; n=19) and subsequent Omicron reinfection (purple; n=13) or vaccination (blue; n=11); ii) initial primary pre-Omicron infection (yellow; n=54), followed by secondary Omicron BA.1/2 infection (orange; n=23); iii) primary vaccination (pink; n=15) followed by breakthrough Omicron infection (black; n=6). Inset percentages indicates the geometric mean BA.1 antibody level in respect to the Wuhan-Hu-1 specific antibody level. Two tailed paired Wilcoxon test, bars indicate geometric mean ±95% CI. B. Ratio of antibody binding against Omicron or ancestral Wuhan-Hu-1 spike (Omicron/ancestral) in children following initial primary Omicron or pre-Omicron infection, or vaccination as A). Kruskal-Wallis test with Dunn’s multiple comparisons test, bars indicate median ±95% CI.

A very different pattern was observed in children who had initial infection with pre-Omicron virus. Here the Omicron:pre-Omicron binding ratio was only 0.14 following primary infection, indicating strong deviation of recognition towards ancestral spike. This increased only modestly to 0.38 following Omicron infection. A similar profile was seen in children following primary vaccination or Omicron breakthrough infection with values of 0.34 in both cases (**Figure 4A,B**).

These data demonstrate that immune imprinting by the primary viral or vaccine exposure determines the profile of the antibody response following subsequent SARS-CoV-2 challenge.

### SARS-CoV-2-specific cellular responses are similar following differential primary SARS-CoV-2 exposure

T cell responses are critical in the control of SARS-CoV-2 but have not yet been assessed in children following Omicron infection. Importantly, spike-specific cellular responses were present in 94% (16/17) of children following primary Omicron infection (**Figure 5A**). T cell responses were also observed in all children following secondary Omicron infection and the magnitude was broadly equivalent (**Figure 5B**). Furthermore, cellular immunity was comparable against peptide pools from either the ancestral Wuhan-Hu-1 or Omicron BA.1 spike protein. Omicron-specific cellular immunity was also assessed in historical samples from 5 children who had been infected with pre-Omicron variant, and again, no difference was seen in the magnitude of ancestral or Omicron-specific response (**Figure 5C**). As such, T cell immunity against Omicron is comparable following infection with either Omicron or pre-Omicron virus. Analysis of vaccinated children showed cellular responses in 92% of donors (17/19) and again the magnitude was comparable to that seen following natural infection (**Figure 5D**).

**Figure 5.**
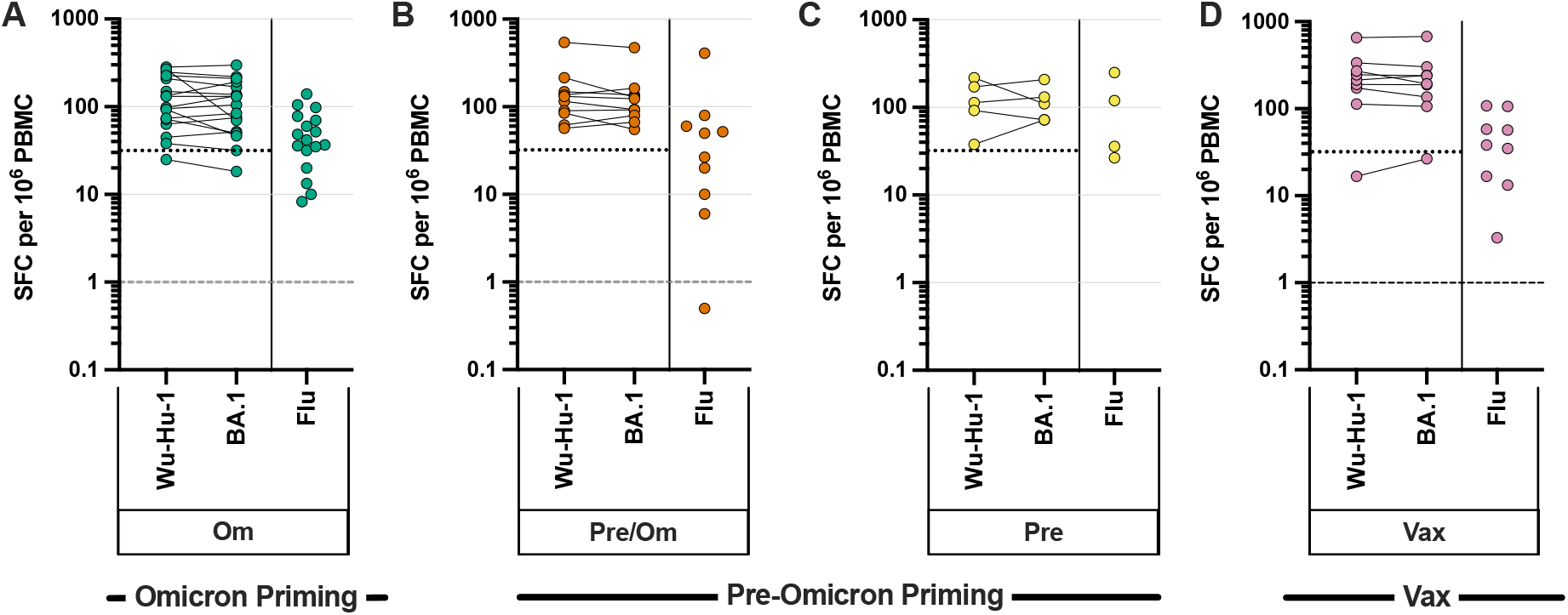
SARS-CoV-2-specific T cell responses are induced reliably following primary Omicron infection and not boosted further by re-infection or vaccination. IFNγ ELISpot was used to assess cellular response to ancestral and Omicron sequence spike peptide pools. Peptides from Influenza (Flu) were included as a control. Children with primary Omicron infection (A, green, n =17), Omicron secondary infection (B, orange, n=10), pre-Omicron infection sampled prior to the Omicron wave (C, yellow, n=5), or vaccinated children (D, pink, n=9). Negative control samples including pre-pandemic paediatric blood samples and DMSO alone incubation were used to define the background level of 1 cell/10^6^ PBMC (16).

These data show that a SARS-CoV-2-specific cellular response develops robustly following primary Omicron infection and is broadly cross-reactive against Omicron and pre-Omicron spike protein. In marked contrast to the humoral response, the magnitude of cellular immunity was not significantly boosted by secondary infection or vaccination.

## Discussion

Omicron SARS-CoV-2 is now an endemic infection and it is important to understand immune responses following infection in children. Our findings highlight a range of unique features regarding Omicron-specific immune responses that have potential implications for the trajectory of the pandemic and future vaccine strategies in this age group.

We initially characterised the immune response following primary Omicron infection where a striking feature was the low magnitude and functional quality of the antibody response. Indeed, a quarter of children failed to demonstrate neutralisation activity against BA.1 despite recent infection, and this concurs with other recent findings (12). These data explain poor protection against reinfection and it was noteworthy that 54% showed serological evidence of re-infection within 3 months, a time period which encompassed emergence of the BA.4/5 subvariants (*5*). A pattern of frequent reinfection is also seen with the four endemic alpha- and beta-HCoV which circulate widely and generally elicit mild upper respiratory tract infection in children (*20*, *21*). Omicron also preferentially replicates within the upper airways and this may underpin a somewhat less severe clinical profile (*22*–*25*) and relatively muted humoral response (*26*).

Although primary HCoV infections elicit modest antibody responses that are unable to prevent recurrent infection (*19*, *21*, *27*, *28*), repeated infections progressively drive maturation of the antibody response which typically achieves adult levels towards the onset of puberty (*19*, *27*, *28*). As such we were interested to assess the immunological impact of Omicron reinfection and found that this enhanced antibody titres and neutralisation activity by 6-fold. We were also able to assess the impact of second antigen challenge through vaccination and the observed marked improvement in humoral immunity against all variants is encouraging for potential future protection (12).

Primary infection with a pre-Omicron virus elicited much stronger humoral immunity against the ancestral virus but notably antibody responses against Omicron spike were comparable to those seen in children after primary Omicron infection. A secondary infection with Omicron in these children delivered a particularly strong antibody boost with impressive neutralisation and exceeded values observed with dual Omicron infection. Pre-Omicron viruses are thus strongly immunogenic for humoral immunity in children and it is interesting to speculate that this may contribute to the development of multisystem inflammatory syndrome (MIS-C) which can be associated with auto-antibody production (*29*).

COVID-19 vaccines are now deployed widely in children and initial vaccination also elicited particularly strong antibody responses. Indeed, a single vaccine dose elicited values that were higher than seen following natural dual infection and this value was doubled following a second dose (Figure 6), revealing potent vaccine immunogenicity in children despite incomplete clinical protection against infection (*5*, *30*).

**Figure 6.**
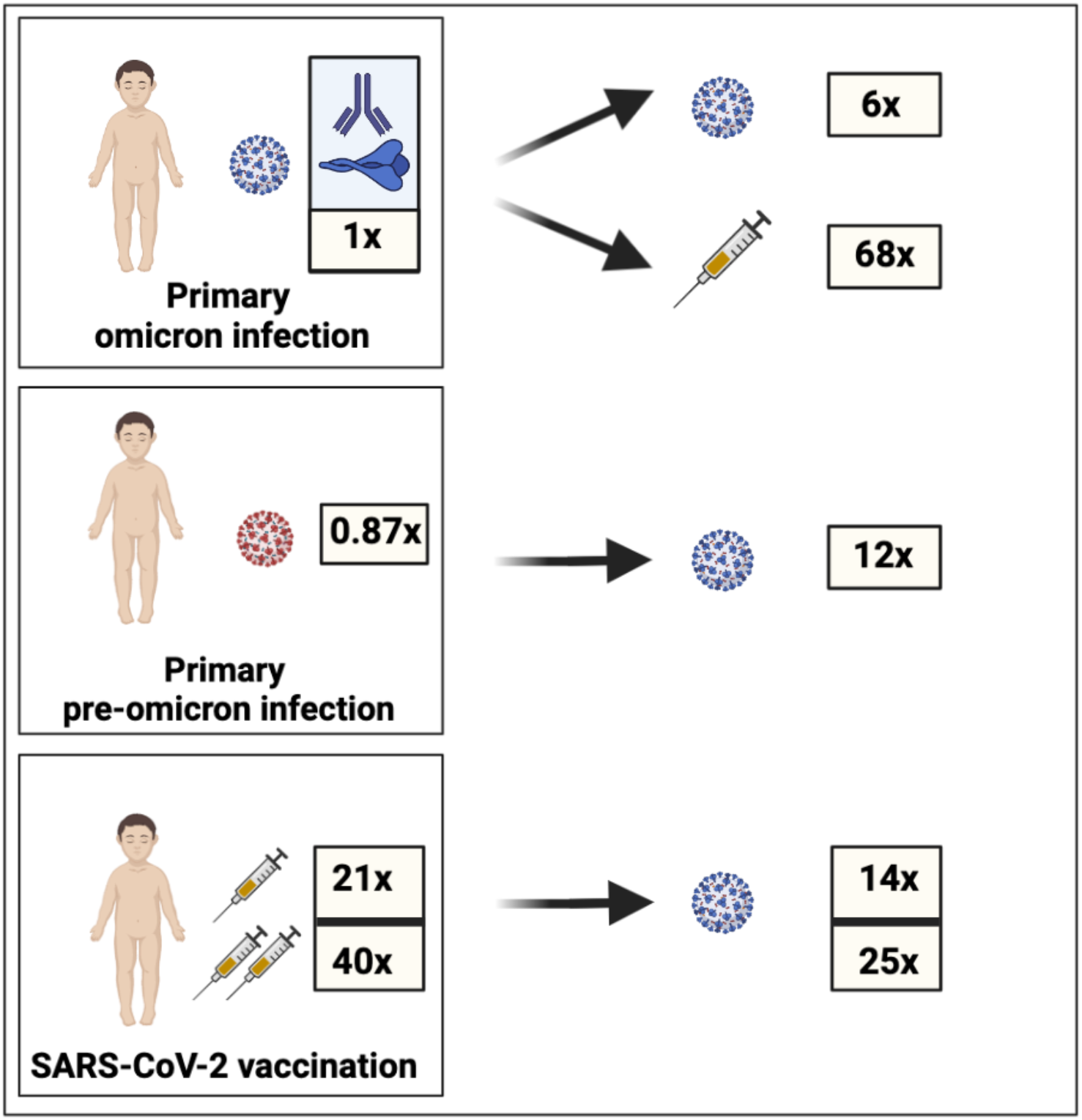
Relative magnitude of omicron-specific antibody response following single or dual SARS-CoV-2 infection or vaccination. Values are given in relation to magnitude following primary Omicron infection. Values for vaccination show response after one or two vaccine doses.

Immune imprinting can limit the capacity to respond to variant antigen challenge (*31*–*33*) and is of considerable concern for long-term protection against SARS-CoV-2 variants. Our prospective assessment of antibody specificity allowed a unique opportunity to address this question and we find that immune imprinting has a profound effect on long term humoral immunity in children. In particular, those who initially experience Omicron infection have a comparable relative antibody response against Omicron and ancestral variants following primary infection and this pattern is maintained following subsequent challenge with either Omicron (reinfection) or ancestral (vaccination) spike. This profile differs somewhat from a predominant Omicron-focussed antibody response reported in adults following primary infection (*11*). In contrast, those who are initially exposed through pre-Omicron infection or vaccine develop humoral immunity deviated markedly towards the ancestral spike protein and this profile also remains broadly stable following challenge with Omicron secondary infection or breakthrough infection after vaccination (**Figure 6**). This profile of ‘antigenic seniority’ is also seen following primary influenza infection and results in lifelong imprinting of immune responses that can impact clinical protection against influenza A subtypes many decades later (*34*, *35*). Our findings show that similar imprinting of antigenic seniority occurs following SARS-CoV-2 infection in children, the future consequence of which is currently unknown.

However, it is reassuring that Omicron-specific antibody and neutralisation titres were increased following Omicron secondary infection or vaccination and this is compatible with recent murine fate mapping experiments which show that *de novo* B cell responses against Omicron could develop due to the degree of antigenic divergence between the variants and that these primary clones targeted novel epitopes within the Omicron domain. We thus provide evidence suggesting that, whilst antigenic seniority is evident, this may not functionally impinge on future protective responses in children.

Cellular immunity is an important mediator of protection against severe COVID-19 (*36*). We find that robust T cell responses develop following either Omicron or pre-Omicron infection. Furthermore, and in direct contrast to humoral immunity, the magnitude of these responses is comparable. This response was also stable after re-infection and similar to that seen following vaccination. As such, cellular immunity appears to be highly sensitive to SARS-CoV-2 infection and may reach a relative plateau following a range of differential initial antigen exposures. Cellular responses against Omicron and ancestral virus were broadly comparable, reflecting the ability of cellular immune recognition to tolerate viral mutations that evade humoral recognition (*37*). T cell responses against endemic HCoV are also robust and somewhat higher in younger people (*38*), further suggesting a differential age-dependent priming of immunity against coronaviruses.

Limitations of the study include absence of samples from vaccinees prior to infection and lack of prospective viral screening to accurately define reinfection following national withdrawal of community and home testing for SARS-CoV-2 after March 2022.

Our results provide important new insights into SARS-CoV-2 immune responses in children and suggest that the Omicron variant of SARS-CoV-2 has evolved to represent a fifth endemic coronavirus infection immunologically. These findings have implications for the future course of the COVID-19 pandemic and its impact on children. Whilst initial Omicron exposure does not reliably generate protective immunity against reinfection this is enhanced following re-exposure and is broadly comparable against Omicron subvariants. Robust cellular responses should support clinical protection against severe disease. Furthermore, the nature of the initial SARS-CoV-2 exposure dictates the subsequent profile of antibody response to antigen challenge and may direct the lifelong pattern of response to SARS-CoV-2 variants and vaccines. Immune imprinting will act as an important individual determinant of humoral immunity, the long-term immunological and clinical importance of which is uncertain.

## Acknowledgments

The authors would like to express their gratitude to the children who took part in the Born in Bradford and sKIDs surveillance studies, and also thank the schools, headteachers, staff, and families of those involved.

## Author Contributions

Conceptualization: ACD, JW, SNL, PM

Methodology: ACD, BW, PM

Investigation: ACD, TL, RB, CB, PS, JZ, SS, AZ, JB, TR, CS, UA, SD, RS

Resources: GI, AAP, AB, BB, FB, IO, JB, SA, FA, JG, MER, DW, JW, SNL

Visualization: ACD, PM

Funding acquisition: BW, JW, SNL, PM

Project administration: GI, AAP, DW

Supervision: ACD, RB, RA, DW, BW, JW, SNL, PM

Writing – original draft: ACD, SNL, PM

Writing – review & editing: All authors

## Methods

### Sample collection

Samples were collected from two prospective studies within the paediatric population.

The SARS-CoV-2 surveillance in primary schools (sKIDs) study of children aged 6-14 years was a cross sectional 5-10ml blood collection study initiated in June 2020 by the United Kingdom Health Security Agency (UK-HSA) after schools reopened following the easing of national lockdown (https://www.gov.uk/guidance/covid-19-paediatric-surveillance). Further collections for this study were undertaken after 21 months (March/April 2022) and 24 months (June/July 2022).

The Born in Bradford study is a prospective longitudinal collection of 5-10ml blood samples from children aged 11-14 years in Bradford, UK undertaken to study SARS-CoV-2 immune responses (*17*). Initial sampling was undertaken at November 2020 and 3 additional samples from donors were collected at 6 monthly intervals until November 2022.

Omicron infection was determined through linkage with the national SARS-CoV-2 testing database (SGSS) held by UKHSA and the NIMS database, which records all COVID-19 vaccinations in England, was used to obtain records of COVID-19 vaccination and vaccine manufacturer for each dose. These data were accessed in May 2022.

No statistical methods were used to predetermine sample sizes. Researchers were blinded to the status of donors before ELISpot and serological assessment.

Ethical review for the sKIDs study was provided by the PHE Research Ethics and Governance Group (PHE R&D REGG ref. no. NR0209). Ethical Review of The Born in Bradford study was provided by the National Health Service Health Research Authority Yorkshire and the Humber (Bradford Leeds) Research Ethics Committee; REC reference: 16/YH/0320. Children and parents or guardians were provided with age-appropriate information sheets prior to enrolment. Written informed consent was obtained from all from parents or guardians of all participants.

### PBMC and Plasma Preparation

Lithium Heparin blood tubes were processed within 24hrs of collection. Briefly tubes were spun at 300g for 10mins prior to removal of plasma which was then spun at 800g for 10mins and stored as aliquots at −80°C. Remaining blood was diluted with RPMI and PBMC isolated on a Sepmate ficol density gradient (Stemcell), cells were washed with RPMI and rested in RPMI+10% batch tested FBS for a minimum of 4 hours prior to cellular assays.

### Serological analysis of SARS-CoV-2-specific immune response

Quantitative IgG antibody titres were measured using Mesoscale Diagnostics multiplex assays; Coronavirus Panel 22, 24, 25 and 27 as previously describe (*15*) following the manufacturer instructions. Briefly, samples were diluted 1:5000 and added wells of the 96 well plate alongside reference standards and controls. After incubation, plates were washed and anti-IgG-Sulfo tagged detection antibody added. Plates were washed and were immediately read using a MESO TM QuickPlex SQ 120 system. Data was generated by Methodological Mind software and analysed with MSD Discovery Workbench (v4.0) software. Data are presented as arbitrary units (AU)/ml determined relative to a 8-point standard curve generated on each MSD plate with standards provided by MSD. Cut-offs were previously defined against pre-pandemic adult and paediatric samples (*15*). Wuhan-Hu-1 RBD-specific antibody titres above or below the geometric mean titre was used to define primary or secondary Omicron infection respectively.

### Pseudotype-based neutralisation assays

Constructs and 293-ACE2 cells were previously described (*15*, *39*). The assay was performed as previously described (*15*, *39*), briefly neutralising activity in each sample was measured against pseudo-virus displaying SARS-CoV-2 spike, either B.1 sequence or B.1.1.529:BA.1, BA.2, BA.4, BA.2.12.1, or BA.2.75 sequence, by a serial dilution approach. Each sample was serially diluted in triplicate from 1:50 to 1:36450 in complete DMEM prior to incubation with approximately 1×10^6^ CPS (counts per second) per well of HIV (SARS-CoV-2) pseudotypes, incubated for 1 hour, and plated onto 239-ACE2 target cells. After 48-72 hours, luciferase activity was quantified by the addition of Steadylite Plus chemiluminescence substrate and analysis on a Perkin Elmer EnSight multimode plate reader (Perkin Elmer, Beaconsfield, UK). Antibody titre was then estimated by interpolating the point at which infectivity had been reduced to 50% of the value for the no serum control samples.

### IFN-γ ELISpot

T cell responses were measured using a IFN-γ ELISpot Pro kit (Mabtech) as previously described. (*15*) Pepmixes pool containing 15-mer peptides overlapping by 10aa from either SARS-CoV-2 spike S1 or S2 domains from the Wuhan-Hu-1 (protein ID: P0DTC2) or Omicron (B1.1.529; BA.1) variant were purchased from JPT technologies. Overlapping pepmixes from influenza A matrix protein 1, California/08/2009 (H1N1) (protein ID: C3W5Z8) and A/Aichi/2/1968 H3N2 (protein ID: Q67157) were purchased from JPT Peptide Technologies and combined as a relevant control.

Briefly, fresh PBMC were rested overnight prior to assay and 0.25-0.3×10^6^ PBMC were added in duplicate per well containing either pep-mix, or DMSO (negative) control, or a single well containing anti-CD3 (positive). Samples were incubated for 16-18hrs. Plates were developed following the manufacturer’s instructions and read using an AID plate reader (AID).

### Data visualisation and statistics

Data was visualised and statistical tests, including normality tests, performed as indicated using GraphPad Prism v9 software. Only results found to be significant (p<0.05) are displayed.

## Extended Data Tables and Figures

**Extended Data Table 1.**
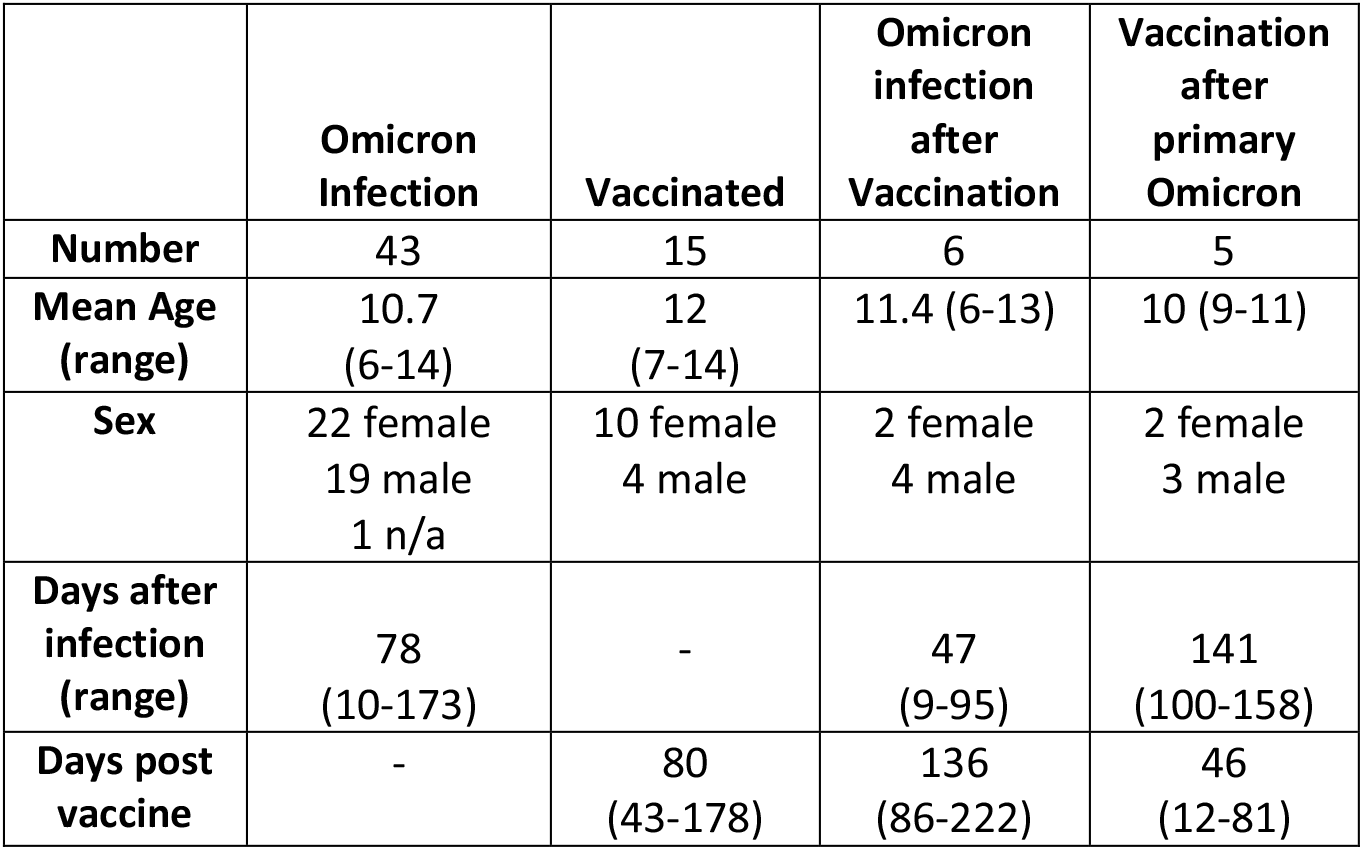
Cohort Demographics

**Extended Data Table 2.** Antibody levels in cohorts. Geometric mean (95% CI of geo. mean).

| Cohort |  | Antibody (AU/ml) |  |  |  |
| --- | --- | --- | --- | --- | --- |
|  |  | Spike-Wuhan<br>-Hu-1 | Spike-Omicron<br>BA.1 | RBD- Wuhan<br>-Hu-1 | RBD- Omicron<br>BA.1 |
| **Pre-Omicron** | **N=54** | 15333<br>(12360-19,020) | 2188<br>(1770-2710) | 4546<br>(3580-5,770) | 538<br>(426-680) |
| **Primary**<br>**BA.1/2**<br>**infection** | **N=20** | 3401<br>(2470-4680) | 2504<br>(1530-4110) | 470<br>(290-770) | 411<br>(245-640) |
| **Secondary**<br>**BA.1/2**<br>**infection** | **N=23** | 82850<br>(59254-115852) | 33187<br>(18530-59440) | 31922<br>(21510-47370) | 7774<br>(4080-14820) |
| **Vaccinated** | **N=15** | 235899<br>(145573-382273) | 81320<br>(40874-161788) | n.d. | n.d. |
| *Vaccinated –*<br>*First dose only* | *N=5* | *207307*<br>*(49961-860196)* | *53710*<br>*(7000-412135)* | n.d. | n.d. |
| *Vaccinated –*<br>*Second dose*<br>*only* | *N=10* | *251642*<br>*(145643-434788)* | *100061*<br>*(47911-208978)* | n.d. | n.d. |
| **Omicron post**<br>**vaccination** | **N=6** | 122202<br>(65947-226445) | 42091<br>(14195-124808) | n.d. | n.d. |
| **Vaccination**<br>**post-primary**<br>**BA.1/2** | **N=11** | 126676<br>(69341-231419) | 169722<br>(52685-546751) | n.d. | n.d. |
| **Secondary**<br>**exposure**<br>**post-primary**<br>**BA.1/2** | **N=13** | 17096<br>(8289-32257) | 15001<br>(6523-34498) | n.d. | n.d. |

**Extended Data Figure 1.**
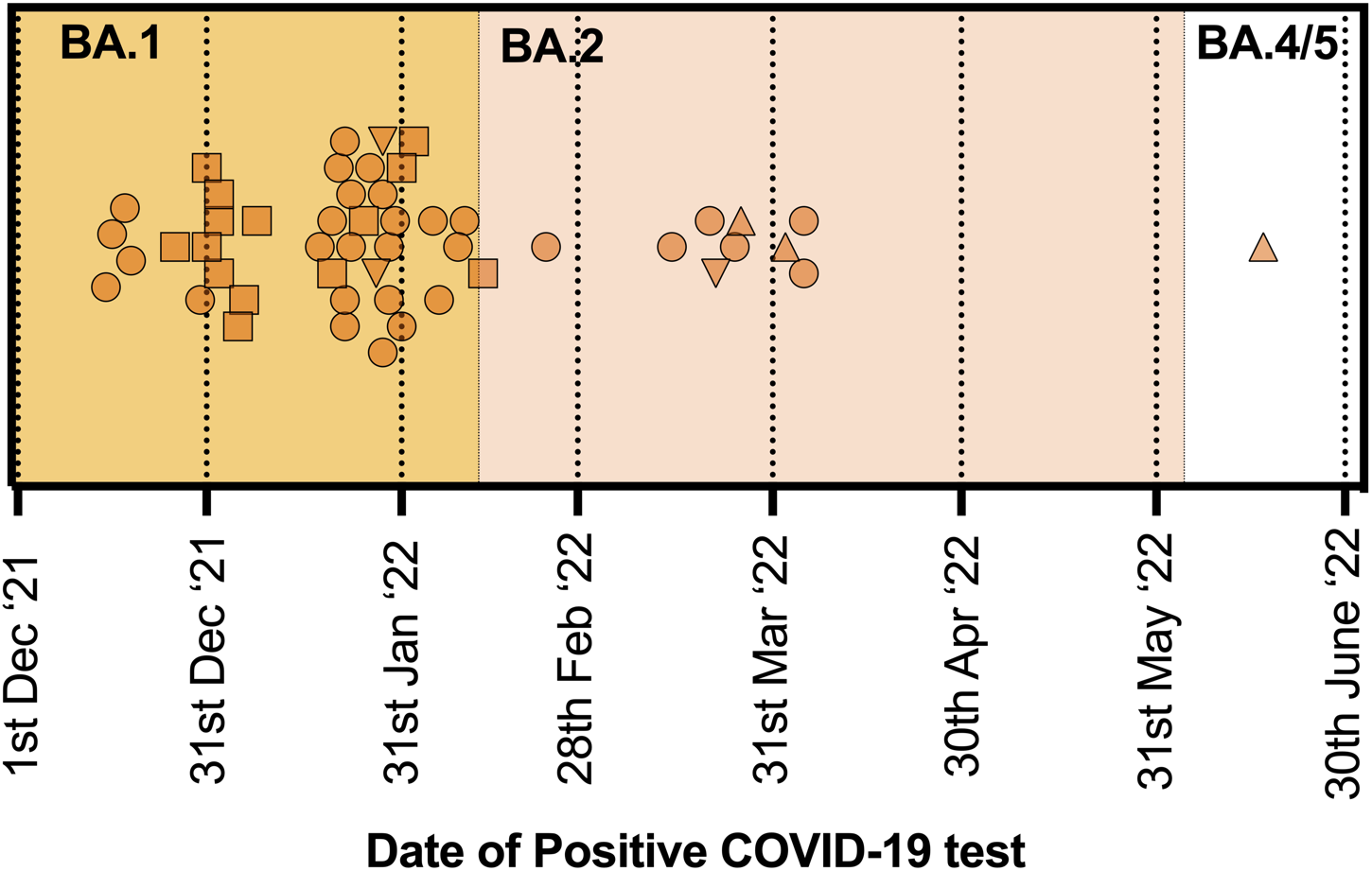
Timing of SARS-CoV-2 Infection in relation to Omicron variant prevalence. A) Timing of positive COVID-19 test results for sKIDs (circles) and Born in Bradford (squares) samples. Triangles indicate breakthrough infections after vaccination. The majority of infections occurred during the BA.1 wave (yellow) whilst the pink shaded area indicates the period in which BA.2 comprised over 50% of sequenced infections nationally. The emergence of BA.4/5 was seen from June 2022 onwards.

**Extended Data Figure 2.**
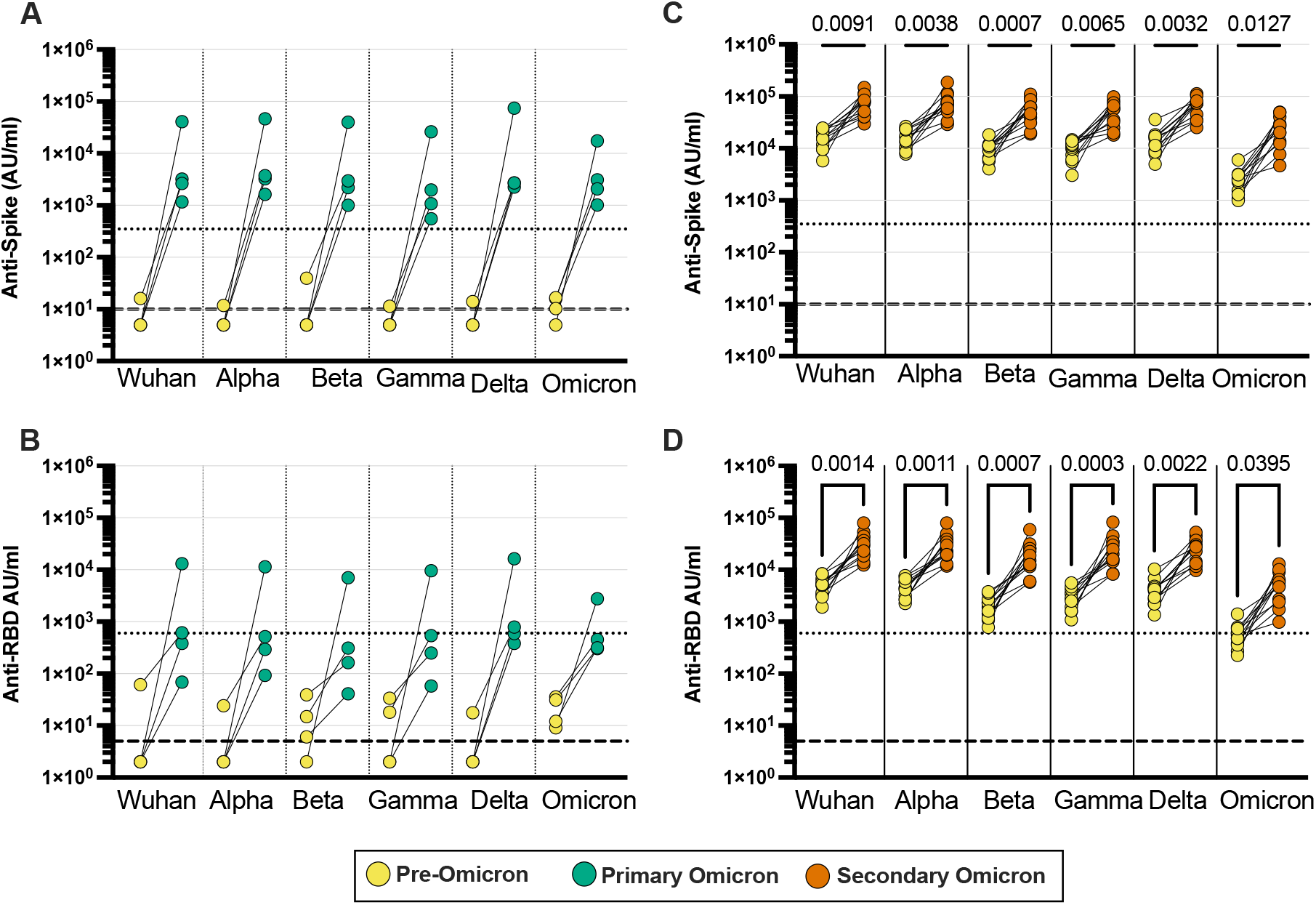
Profile of SARS-CoV-2-specific antibody response following primary or secondary Omicron infection. SARS-CoV-2-specific antibody responses were determined in longitudinal plasma samples from children with BA.1/2 infection. Spike (A) and RBD-specific (B) antibody binding against viral variants in samples from children (n=4) who were seronegative prior to Omicron infection (yellow) and following BA.1/2 infection (green). Spike (C) and RBD-specific (D) antibody binding against viral variants in samples from children (n=11) who were seropositive prior to BA.1/2 infection (yellow) and following BA.1/2 infection (orange). Dotted lines indicate sero-positive cut-offs as determined for Wuhan-Hu-1-specific antibody response; dashed lines indicate below limit of detection. Friedman Test with Dunn’s multiple comparisons test.

**Extended Data Figure 3.**
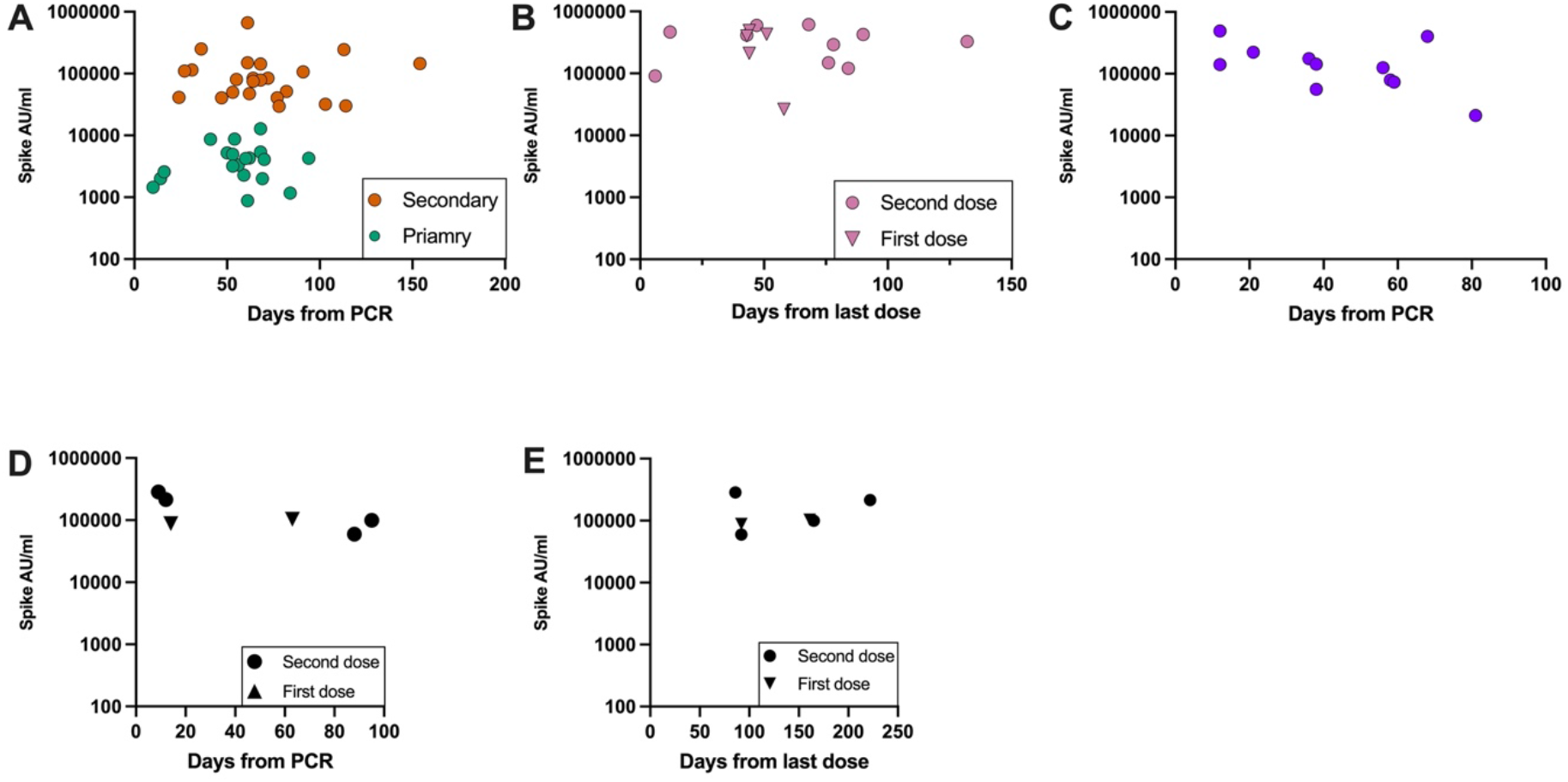
Antibody titre in respect of time from infection or vaccination. The prospective nature of sample collection was such that children were studied at varying timepoints after infection or vaccination. Antibody titre against ancestral B.1 spike was therefore plotted against time since antigen challenge to assess the potential impact of antibody waning. A) Time from primary (green) or secondary (orange) Omicron infection. B) Time from a first (triangle) or second (circle) COVID-19 vaccine dose. C) Time from single vaccine dose in children with prior Omicron infection. D) Time from Breakthrough infection or (E) last vaccine dose in vaccinated children who experienced a breakthrough Omicron infection.

**Extended Data Figure 4.**
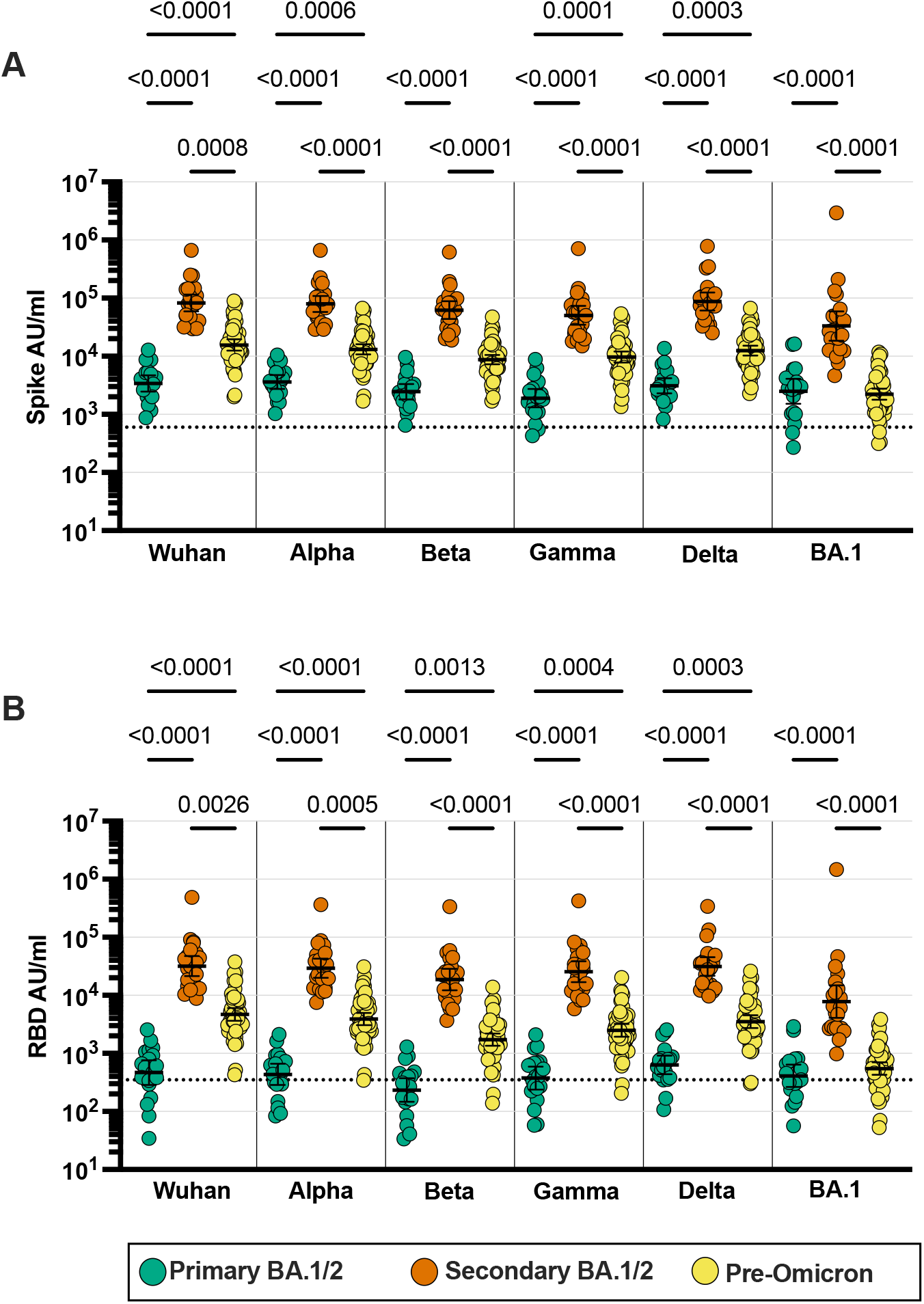
Relative antibody titre against SARS-CoV-2 viral variants following primary or secondary Omicron infection. Samples from children (n=43, aged 6-14 years) recently infected with Omicron BA.1/2 variant were assessed on the MSD-platform for antibodies against Spike protein (A) and RBD-domain (B) from SARS-CoV-2 variants as indicated. Donors were divided in those with primary SARS-CoV-2 infection (green, n=20) or secondary SARS-CoV-2 infection (Orange, n=23). Values are also compared to antibody levels from historical children’s samples taken after SARS-CoV-2 infection prior to emergence of Omicron (yellow, n=54, aged 5-14 years). Friedman Test with Dunn’s multiple comparisons test.

**Extended Data Figure 5.**
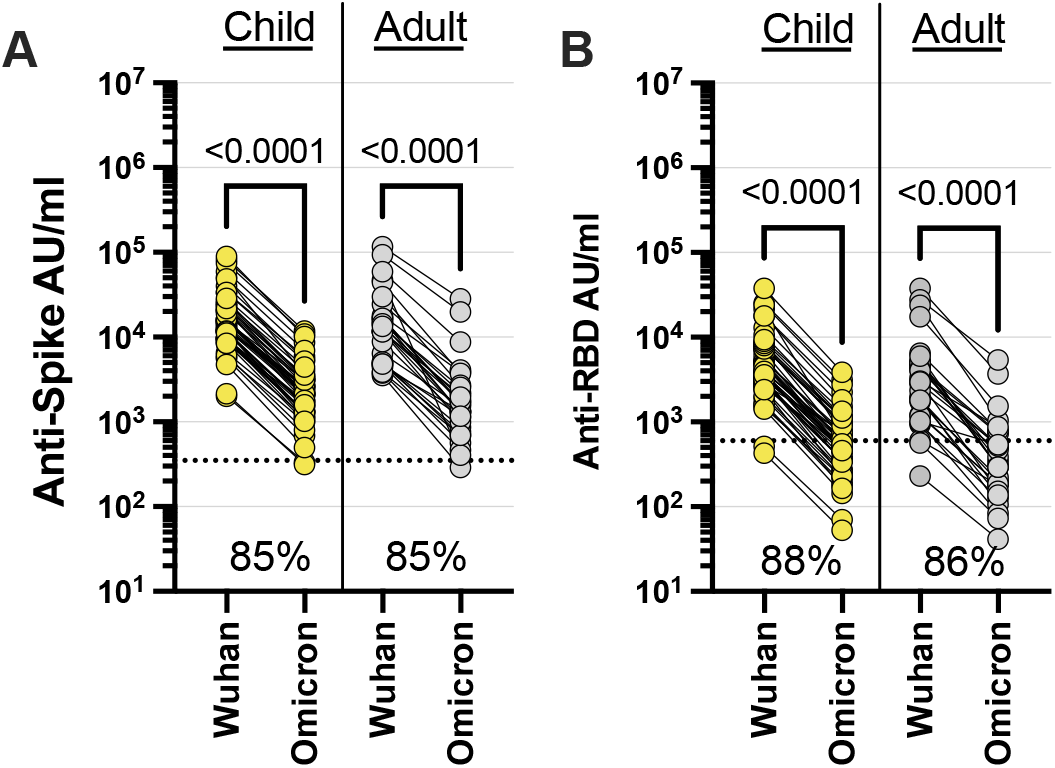
Marked reduction in Omicron-specific antibody binding in sera from children with pre-Omicron natural infection. Antibody binding to Wuhan-Hu-1 or BA.1 spike (A) or RBD-domain (B) in seropositive children (n=54, aged 5-14 years) and adults (n=30; aged >18 years), infected and sampled prior to the emergence of Omicron. Lines join individual donors. Inset percentage indicates the average reduction in binding (AU/ml) to Omicron protein compared to Wuhan. Kruskal-Wallis test with Dunn’s multiple comparisons test. Dotted lines indicate seropositive cut-offs as determined for Wuhan-specific response.

